# Neurotoxic Amyloidogenic Peptides Identified in the Proteome of SARS-COV2: Potential Implications for Neurological Symptoms in COVID-19

**DOI:** 10.1101/2021.11.24.469537

**Authors:** Saba Islam, Mirren Charnley, Guneet Bindra, Julian Ratcliffe, Jiangtao Zhou, Raffaele Mezzenga, Mark Hulett, Kyunghoon Han, Joshua T. Berryman, Nicholas P. Reynolds

## Abstract

COVID-19 is primarily known as a respiratory disease caused by the virus SARS-CoV-2. However, neurological symptoms such as memory loss, sensory confusion, cognitive and psychiatric issues, severe headaches, and even stroke are reported in as many as 30% of cases and can persist even after the infection is over (so-called ‘long COVID’). These neurological symptoms are thought to be caused by brain inflammation, triggered by the virus infecting the central nervous system of COVID-19 patients, however we still don’t fully understand the mechanisms for these symptoms. The neurological effects of COVID-19 share many similarities to neurodegenerative diseases such as Alzheimer’s and Parkinson’s in which the presence of cytotoxic protein-based amyloid aggregates is a common etiological feature. Following the hypothesis that some neurological symptoms of COVID-19 may also follow an amyloid etiology we performed a bioinformatic scan of the SARS-CoV-2 proteome, detecting peptide fragments that were predicted to be highly amyloidogenic. We selected two of these peptides and discovered that they do rapidly self-assemble into amyloid. Furthermore, these amyloid assemblies were shown to be highly toxic to a neuronal cell line. We introduce and support the idea that cytotoxic amyloid aggregates of SARS-CoV-2 proteins are causing some of the neurological symptoms commonly found in COVID-19 and contributing to long COVID, especially those symptoms which are novel to long COVID in contrast to other post-viral syndromes.

## Introduction

The disease caused by viral infection with SARS-COV-2 is known as COVID-19 and whilst predominantly a respiratory disease affecting the lungs it has been shown to display a remarkably diverse array of symptoms. These include a range of moderate to severe neurological symptoms that persist for up to 6 months after infection reported in as many as 30% of patients.^1^ These symptoms include memory loss, sensory confusion (such that previously pleasant smells become fixed as unpleasant, for example) cognitive and psychiatric issues, severe headaches, brain inflammation and haemorrhagic stroke.^1–5^ The molecular origin of these neurological symptoms is as yet unknown but they are similar to hallmarks of amyloid related neurodegenerative diseases such as Alzheimer’s (AD),^6, 7^ and Parkinson’s.^8^ Impaired olfactory identification ability in particular has been reported in AD and prodromal AD ailments, such as mild cognitive impairment (MCI).^9^ COVID-19 related anosmia and phantosmia has been shown to correlate with a persistence of virus transcripts in the olfactory mucosa and in the olfactory bulb of the brain, and with persistent inflammation, however negative evidence for continuing viral replication has also been shown for long-term anosmia.^10^

Proteins from the Zika virus^11^ and also the coronavirus responsible for the Severe Acute Respiratory Syndrome (SARS) outbreak in 2012 (Sars-CoV-1)^12^ have been shown to contain sequences that have a strong tendency to form amyloid nanofibrils. As the proteome of SARS-COV-1 and SARS-COV-2 possess many similarities^13, 14^ we propose amyloid nanofibrils formed from proteins in SARS-COV-2 may be responsible for the neurological symptoms in COVID-19. Further, there is evidence that SARS-COV-2 is neuroinvasive with either the full virus^15, 16^ or viral proteins^16^ being found in the CNS of mouse models and the post-mortem brain tissue of COVID-19 patients. Therefore, amyloid forming proteins from the SARS-COV-2 virus in the CNS of COVID-19 infected patients could have similar cytotoxic and inflammatory functions to amyloid assemblies that are the molecular hallmarks of amyloid related neurodegenerative diseases such as Alzheimer’s (A*β*, Tau) and Parkinson’s (*α*-synuclein).

Viral genomes evolve rapidly and are highly constrained by size, therefore every component is typically functional either to help the virus replicate or to impede the host immune system, allowing also some constraints to genome physical structure. If the proteome of SARS-COV-2 does contain amyloid forming sequences, this raises the question “what is their function?” To this end, there are a number of potential roles for amyloid assemblies in pathogens generally^17^ and specifically in coronaviruses such as SARS-COV-2. The simplest is that amyloid is an inflammatory stimulus^18^, and pro-inflammatory cytokines can up-regulate expression of the spike protein receptor ACE-2 such that intercellular transmissibility of SARS-COV-2 is increased^19^.

Eisenberg and co-workers found that the Nucleocapsid protein (NCAP) in SARS-COV-2, which is responsible for packaging RNA into the virion, contains a number of highly amyloidogenic short peptide sequences within its intrinsically disordered regions (IDRs).^20^ It has also been shown that the self-assembly of these peptides is enhanced in the presence of viral RNA, during liquid-liquid phase separation (LLPS is an important stage in the viral replication cycle).^21, 22^ These findings suggest amyloids may play an important role in RNA binding and packaging during the viral replication cycle.

It is also possible that amyloid assemblies in coronaviruses might have a role in inhibiting the action of the host antiviral response. Mitochondrial antiviral-signalling proteins (MAVS) are activated during viral infections, this activation is a binary effect where nearly all the MAVS in a cell are activated in quick succession, this triggers the cell to process antiviral molecules such as type I interferons and pro-inflammatory cytokines.^23^ It is believed that this rapid activation occurs by a catalytic prion-like process where the MAVS self-assemble into fibrillar structures on the surface of the mitochondria. Amyloids formed by the SARS-COV-2 proteome may interact with the prion-like MAVS, binding to and inhibiting their self-assembly. This would prevent the binary activation of the MAVS and inhibit the antiviral signalling effects. Such an inhibitory mechanism based on co-assembly or amyloid self-recognition would closely resemble action of peptide inhibitors which have been developed as potential anti-amyloid therapies for Alzheimer’s and Parkinson’s diseases.^24^ Similar co-assembly mechanisms between murine cytomegaloviruses and human RIPK3 kinase have been seen by Pham et. al.^25^ These hybrid amyloids are also proposed to inhibit the signalling capabilities of the host protein.

In a related mechanism to the inhibition of the prion-like activity of the MAVS it is possible that amyloids within SARS-COV-2 might sabotage the antiviral action of lipid droplets. Such lipid droplets have been found to play vital roles in controlling the antiviral immune response through modulation of type-1 interferon and viral replication rates in a number of different viral infections including COVID-19.^26–29^ Furthermore, the importance of lipidic structures (extracellular vesicles, lipid droplets etc) in the antiviral response is becoming more apparent.^30, 31^ Therefore, as amyloids are well known to be efficient binders and disrupters of lipid membranes^32–35^, it is entirely possible that amyloids in viral proteomes may have evolved to disrupt the membrane of antiviral lipid bodies either destroying them totally, or reducing their antiviral properties.

In this study we choose to focus on a selection of proteins from the SARS-COV-2 proteome known as the open reading frames (ORFs). These ORF proteins were chosen as they have no obvious roles in viral replication^36^, perhaps freeing them up to have yet uncharacterised roles in disrupting the host antiviral responses. By sequence and length they appear to be largely unstructured, making them good candidates for amyloid formation *in vivo*. We performed a bioinformatic screening of the ORF proteins to look for potential amyloidogenic peptide sequences. This analysis was used to select two subsequences, one each from ORF6 and ORF10, for synthesis. The synthesised peptides were both found to rapidly self-assemble into amyloid assemblies with crystal, needle and ribbonlike morphologies. Cytotoxicity assays on neuronal cell lines showed these peptide assemblies to be highly toxic at concentrations as low as 0.0005%.

Since commencing this work, others have found that ORF6 is the most cytotoxic single protein of the SARS-CoV-2 genome, showing localisation to membranes when overexpressed in human and primate immune cell lines.^37^ In contrast, ORF10 has been reported as an unimportant gene with very low expression and no essential role in virus replication,^38^ however the functions of immune suppression, inflammation promotion, or membrane disruption via amyloid formation would be non-essential, if present, and should not necessarily require transcription in large volumes, making ORF10 an intriguing second candidate for the present study. ORF8 and ORF10 are the only two coded proteins present in SARS-CoV-2 which do not have a homologue in SARS-CoV.^39^ While long-term consequences from SARS-CoV were severe, including tiredness, depression, and impaired respiration, few or zero unequivocally neurological post-viral symptoms were recorded from the (admittedly quite small) set of documented cases.^40^

The worst-case scenario given the present observations is that of progressive neurological amyloid disease being triggered by COVID-19. To the authors’ knowledge there has so far been no discussion or any documented example of this, however it has been noted that up-regulation of Serum amyloid A protein driven by inflammation in COVID-19 seems like a probable trigger for the the systemic (non-neurological) amyloid disease AA amyloidosis,^41^ which is already known to be a concomitant of inflammatory disease in general.

## Materials & Methods

### Amyloid Prediction Algorithms

The online servers providing TANGO and ZIPPER were used to predict peptide sequences with a tendency to form *β*-rich amyloid assemblies. TANGO is an algorithm that predicts aggregation nucleating regions in unfolded polypeptide chains.^42^ It works on the assumption that the aggregating regions are buried in the hydrophobic core of the natively folded protein. ZIPPER is an algorithm that predicts hexapeptides within larger polypeptide sequences that have a strong energetic drive to form the two complementary *β*-sheets (known as a steric zipper) that give rise to the spine of an amyloid fibril.^43^ Both methods are physically motivated but rely on statistically determined potentials.

### Self-Assembly of Peptides

NH2-ILLIIM-CO2H and Ac-RNYIAQVD-NH2 (> 95% pure) were purchased from GL Biochem Ltd (Shanghai, China). Ideally it would have been preferred to have both peptides capped (N-terminus: Acetyl and C-terminus: Amide), as they would better represent small fragments of a larger peptide sequence. Due to the fact ILLIIM contains no charged sidechains, synthesising capped sequences to high purity would have been very challenging, therefore only the RNYIAQVD sequence remained capped and the ILLIIM sequence had regular carboxyl and amino termini. Self-assembly was initiated by dissolving the peptides at 1 or 5 mg mL^−1^ in hot PBS (80°C); the peptide solutions were vortexed vigorously and held at 80°C for three hours to ensure maximum dissolution. After a second round of vortexing the peptide suspensions were cooled slowly. Complete assembly was seen after 2 hours.

### Atomic Force Microscopy (AFM)

AFM imaging was performed on a Bruker Multimode 8 AFM and a Nanoscope V controller. Tapping mode imaging was used throughout, with antimony (n)-doped silicon cantilevers having approximate resonant frequencies of 525 or 150 kHz and spring constants of either 200 or 5 Nm^−1^ (RTESPA-525, Bruker or RTESPA-150, Bruker respectively). No significant differences were observed between cantilevers. 50 μl aliquots of the peptide (either at 1 or 5 mg mL^−1^) were drop cast onto freshly cleaved muscovite mica disks (10 mm diameters) and incubated for 20 mins before gently rinsing in MQ water and drying under a nitrogen stream. All images were flattened using the first order flattening algorithm in the nanoscope analysis software and no other image processing occurred. Statistical analysis of the AFM images was performed using the open-source software FiberApp^44^ from datasets of no less than 700 fibers.

### Transmission Electron Microscopy (TEM)

Copper TEM grids with a formvar-carbon support film (GSCU300CC-50, ProSciTech, Qld, Australia) were glow discharged for 60 seconds in an Emitech k950x with k350 attachment. Five *μ*l drops of sample suspension was pipetted onto each grid, allowed to adsorb for at least 30 seconds and blotted with filter paper. Two drops of 2% uranyl acetate were used to negatively stain the particles blotting after 10 seconds each. Grids were then allowed to dry before imaging. Grids were imaged using a Joel JEM-2100 (JEOL (Australasia) Pty Ltd) transmission electron microscope equipped with a Gatan Orius SC 200 CCD camera (Scitek Australia).

### Small and Wide Angle X-ray Scattering (SAXS/WAXS)

SAXS/WAXS experiments were performed at room temperature on the SAXS/WAXS beamline at the Australian Synchrotron. Peptide assemblies in PBS prepared at both 1 and 5 mg mL^−1^ were loaded into a 96-well plate held on a robotically controlled x-y stage and transferred to the beamline via a quartz capillary connected to a syringe pump. Data from the 5 mg mL^−1^ assemblies was discarded due to sedimentation of the assemblies preventing reliable sample transfer into the capillaries. The experiments used a beam wavelength of *λ* = 1.03320 Å^−1^ (12.0 keV) with dimensions of 300 *μ*m × 200 *μ*m and a typical flux of 1.2 ×10^13^ photons per second. 2D diffraction images were collected on a Pilatus 1M detector (SAXS). SAXS experiments were performed at q-ranges between 0.02 and 0.25 Å^−1^ and WAXS experiments were performed at a q-range between 0.1 and 2 Å^−1^. Spectra were recorded under flow (0.15 mL min^−1^) to prevent X-ray damage from the beam. Multiples of approximately 15 spectra were recorded for each time point (exposure time = 1 s) and averaged spectra are shown after background subtraction against PBS in the same capillary.

### Circular Dichroism Spectroscopy

CD spectroscopy was performed using an AVIV 410-SF CD spectrometer. Wavelength spectra were collected between 190 and 260 nm in PBS using 1 mm quartz cuvettes with a step size of 0.5 nm and 2 s averaging time. Data was analysed using the BeStSel method of secondary structure determination.^45^

### Atomistic Modelling

Atomistic models were constructed using the Nucleic Acid Builder^46^. Simulations were run in explicit water (TIP3P^47^) using the ff15ipq forcefield^48^ and the pmemd time integrator^49^. In order to hold the unit cell geometry to values consistent with the observed scattering, the alpha carbon of the central residue of each chain was subjected to a restraining force with spring constant 2 kcal mol^−1^ Å^−2^. The system state after 10 ns of equilibration was stripped of water molecules more than 10 Åfrom any non-hydrogen solute atom, and passed to CRYSOL3 for calculation of orientationally averaged scattering profile given the example state (including the ordered waters from the explicit solvent shell, and also including an approximate treatment of ordered water beyond this shell)^50, 51^.

### Thioflavin T Amyloid Kinetic Assays

Peptide assemblies were made up to concentrations of 1 or 5 mg mL^−1^ suspensions containing 25 *μ*M ThT in PBS. The first fluorescence measurement (t = 0) was recorded immediately after sample preparation. All the samples were then stored at room temperature and fluorescence intensity was recorded at different time points. Measurements were performed in triplicate using a ClarioStar fluorimeter equipped with a 96 well plate reader (excitation wavelength: 440 nm, emission wavelength: 482 nm). A representative curve is shown in Figure 4.

### Cell line and cultures

Human derived neuroblastoma cells (SH-SY5Y) were cultured in DMEM-F12 (Invitrogen) medium supplemented with 10% (v/v) fetal calf serum (FCS), 100 U/mL penicillin and 100 *μ*g/mL streptomycin (Invitrogen, Carlsbad, CA). Cells were cultured at 37°C in a humidified atmosphere containing 5% CO2.

### Cell viability assay

Peptides were serially diluted into DMEM-F12 media and seeded onto SH-SY5Y cells cultured in a 96-well plate. Initial plating cell density was pre-optimised to avoid confluency at endpoint. After 48 h, cell viability was determined using 3-(4,5-dimethylthiazol-2-yl)-2,5-diphenyltetrazolium bromide (MTT) (Sigma-Aldrich, MO) as described previously.^52^ Equivalent MTT assays were performed on cells cultured in the same ratios of PBS to media, but in the absence of peptide assemblies to confirm that the culture conditions were non-toxic (figure S10). Absorbance readings of untreated control wells in 100% cell culture media was designated as 100% cell viability. Statistical analysis was performed by one way ANOVA tests with Tukey comparison in the software GraphPad (Prism). *** = P<0.0001

## Results and Discussion

### Amyloid aggregation prediction algorithms identified two short peptides from ORF6 and ORF10 that are likely to form amyloids

Figure 1 shows selected output from bioinformatics tools applied to predict the amyloidogenicity of peptide sequences within larger polypeptides. Application of the ZIPPER tool to ORF-6 provides more than 10 choices of six-residue windows of the sequence predicted to be highly amyloidogenic (1a), while ORF10 shows only 3 such highly amyloidogenic sequence windows (1b). In order to narrow down our search for candidate peptides we also used the TANGO algorithm, which operates in a similar manner. For ORF6 there are two regions that are predicted to be highly aggregation prone, I_14_LLIIMR and D_30_YIINLIIKNL. The region I_14_-R_20_ overlaps almost perfectly with the hexapeptide I_14_LLIIM identified by ZIPPER. The region 30-40 also contains multiple hits in ZIPPER, but as this study was limited to two candidate peptides we chose ILLIIM as our first candidate as it closely resembles the sequence ILQINS from Hen Egg White Lysozyme which has previously been seen to be highly amyloidogenic (the mutation TFQINS in human lysozyme is disease-linked).^53–55^ Looking now at the TANGO plots for ORF-10 the main aggregation prone sequence is residues F_11_TIYSLLLC, although there are no high stability hexapeptides in this sequence predicted by ZIPPER. The octapeptide R_24_NYIAQVD was chosen due to its zwitterionic residue pair R-D which physically should strongly enhance interpeptide association, despite being too far apart in the sequence to trigger the highly local bioinformatics algorithms. Encouragingly ZIPPER also predicts the hexapeptide NYIAQV contained within RNYIAQVD to be highly amyloidogenic. Based on the outputs from ZIPPER and TANGO and also on experience of making and studying amyloid we selected RNYIAQVD and ILLIIM to be synthesised and their amyloid forming capability investigated.

**Figure 1.**
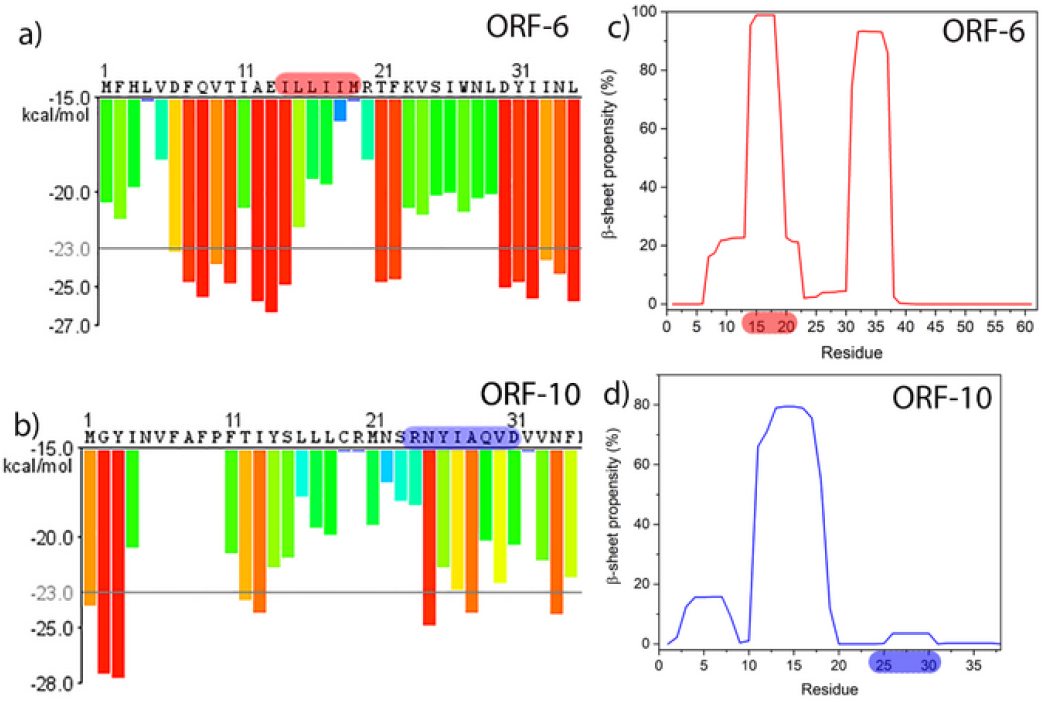
Output from amyloid assembly prediction software for SARS-COV-2 ORF6 and ORF10 sequences. **a-b**) outputs from ZIPPER identifying hexapeptide fragments predicted to form steric zippers nucleating the assembly of amyloid fibrils. Hexapeptides with Rosetta energies below −23 kcal mol^−1^ are predicted to be highly amyloidogenic and are shown with red bars. **c-d**) outputs from TANGO indicating sequence regions of high propensity to form *β*-sheet.

### Nanoscale imaging reveals both peptide sequences self-assembling into fibrillar structures

AFM and TEM imaging of the two peptide assemblies revealed that both peptides assembled in as little as 2 hours at 1 mg mL^−1^ (figures S1 and S2) into needle-like crystalline assemblies. AFM imaging of assemblies formed at either 1 or 5 mg mL^−1^ for 24 h revealed that assemblies from both peptides tend to stack on top of each other forming multilaminar nanofibrillar structures (Figure 2). Evidence of lateral assembly of the needles was also observed but this appears to happen more frequently in the RNYIAQVD assemblies compared to ILLIIM. ILLIIM tends to form very large (2-3 microns in width) multi laminar crystalline assemblies (Figure 2e). AFM was used to investigate the height of the individual assemblies of both ILLIIM and RNYIAQVD. Figure S3 shows a line section through a multilaminar ILLIIM assembly and shows step heights that vary between 4-9 nm, in figure S4 step height analysis of what appears to be a single assembly (no evidence of stacking) appears to be over 12 nm tall. RNYIAQVD tells a similar story, with individual step heights reading around 5.5 nm (Figure S5), but other assemblies having heights of 20 nm or more (figure S6).

**Figure 2.**
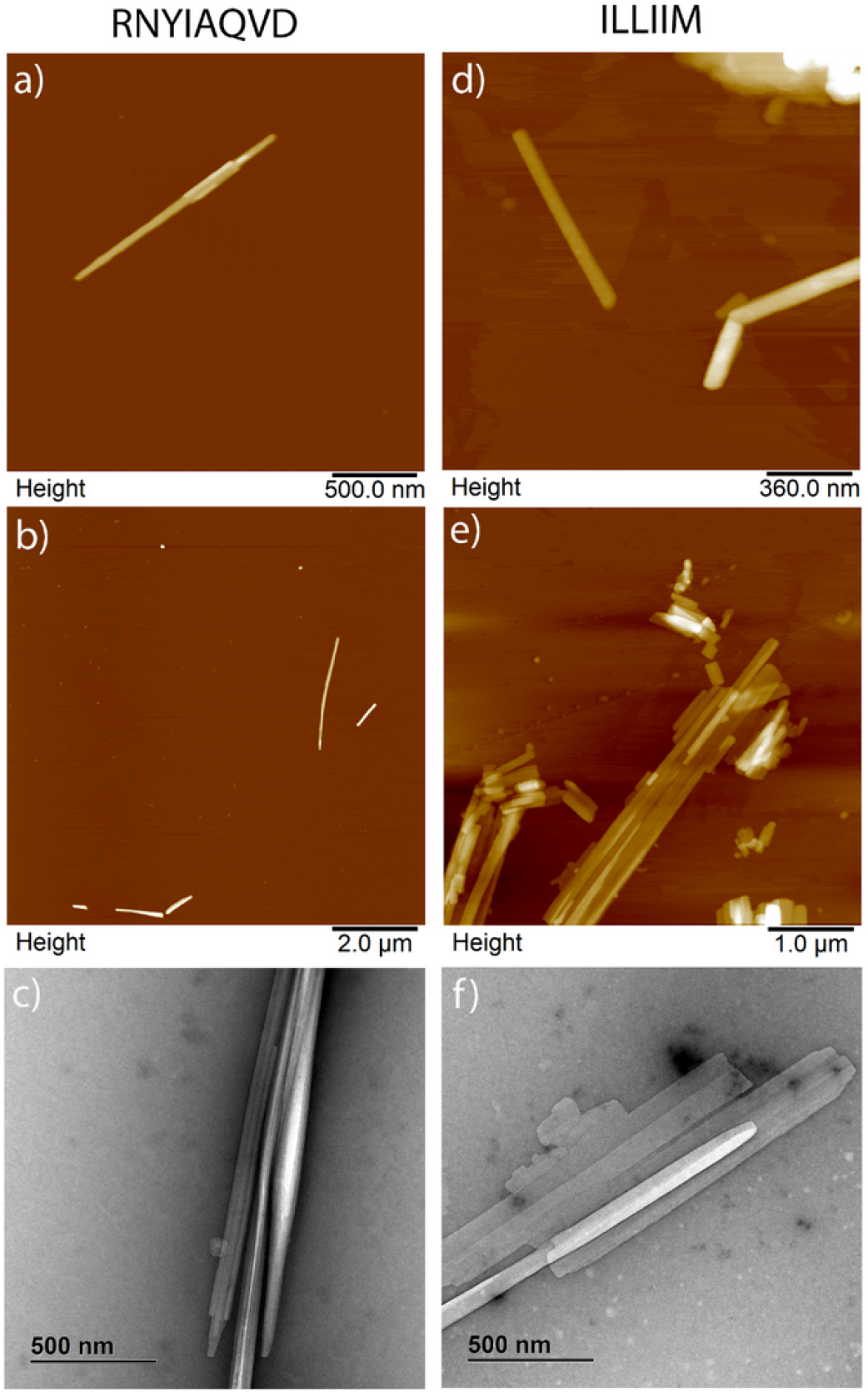
Atomic Force and Transmission Electron Microscopy images of peptide assemblies at 5 mg mL^−1^ incubated for 24 h. a,b) AFM images and c) TEM images of RNYIAQVD assemblies, d,e) AFM image and f) TEM images of ILLIIM assemblies.

Statistical analysis of the fibril widths and contour lengths was performed using the freely available software tool FiberApp.^44^ The analysis of fibril width distribution was taken from the z-axis (height) of the AFM images, both peptide assemblies show a heterogenous distribution of fibril widths due to the previously observed tendency of both assemblies (figure 2 and S1-4) to form multilaminar stacks. Analysis of the distribution of the contour length of the two assemblies showed a biphasic distribution of lengths for both fibrils with two broad sub-populations centred around 1 and 3 *μ*m (figure 3c and 3d) for both peptides. The sub-population at 3 *μ*m was seen to be much larger for the RNYIAQVD peptide (figure 3d) compared to the ILLIIM (figure 3c). This population of longer fibrils correlates with the observation from figure2 that for RNYIAQVD longer, thinner assemblies are favoured (self-assembly via fibril extension) over the wider shorter assemblies more commonly seen for ILLIIM (assembly via lateral association of protofilaments). Analysis of the persistence length of the fibrils (figure S9) showed that whilst both peptides formed very straight linear fibrils, the stiffness of RNYIAQVD (*λ* =41.92 *μ*m) is greater than that of ILLIIM (*λ* = 31.96 *μ* m).

**Figure 3.**
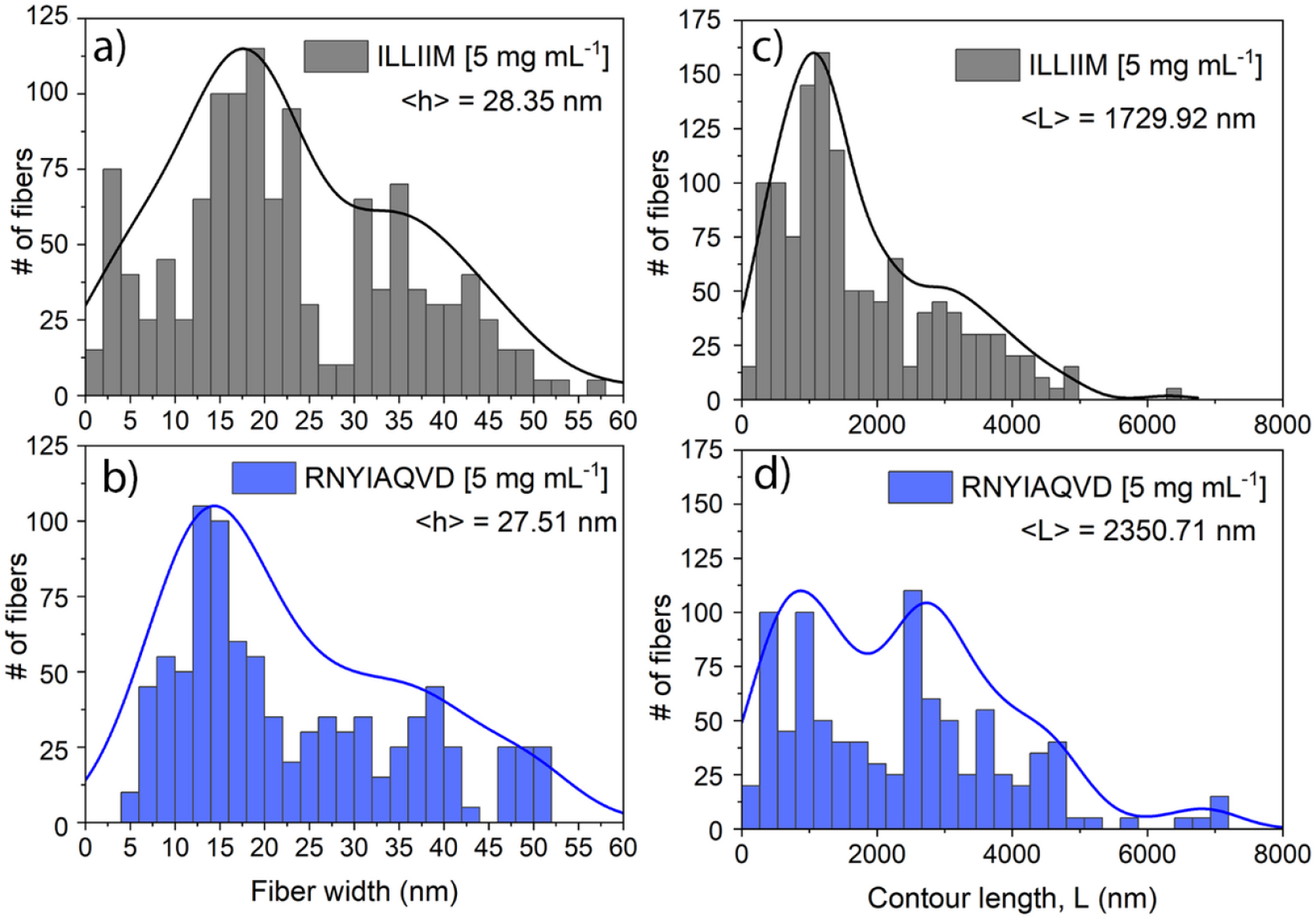
Histograms showing distributions of fiber widths and contour lengths, as determined using the statistical fibril analysis software FiberApp. At least 1000 fibers analysed.

### X-ray scattering, spectroscopic characterisation and molecular modelling confirm the amyloid nature of the assemblies

Figure 4 shows the radially averaged 1D SAXS plots for ILLIIM and RNYIAQVD at the lower concentration studied (at the higher concentration, sedimentation made recording X-ray scattering spectra impossible). In the central part of the scattering curve the ILLIIM assemblies produced a slope with a q^−2^ dependence which is consistent with the form factor of an infinite 2D surface.^53^ Together with the AFM and TEM data this implies a wide flat ribbon architecture similar to that seen in related amyloidogenic short peptides.^53, 56^ RNYIAQVD however, displays a q^−4^ dependence in the (later) central part of the scattering curve, appearing more towards the high-q limit. Porod’s law indicates that q^−4^ scaling (at high q but still less than 0.1Å^−1^) is consistent with any aggregates having sharp and well-defined flat (2D) surfaces but does not otherwise specify shape^57^.

Figure 4b shows the CD spectra of mature assemblies (> 5 days), of both peptides. Assemblies of ILLIIM display a quite simple spectrum indicating dominance of *β*-sheets, with a minimum between 225-230 nm and a strong maximum at 205 nm.^53^ The CD spectra of RNYIAQVD are less obvious and seem to resemble the typical spectra of a random polypeptide coil, except in that there is a well-defined minimum at 203 nm (a typical random coil is < 200 nm) and additional shoulder appears at around 215 nm.

**Figure 4.**
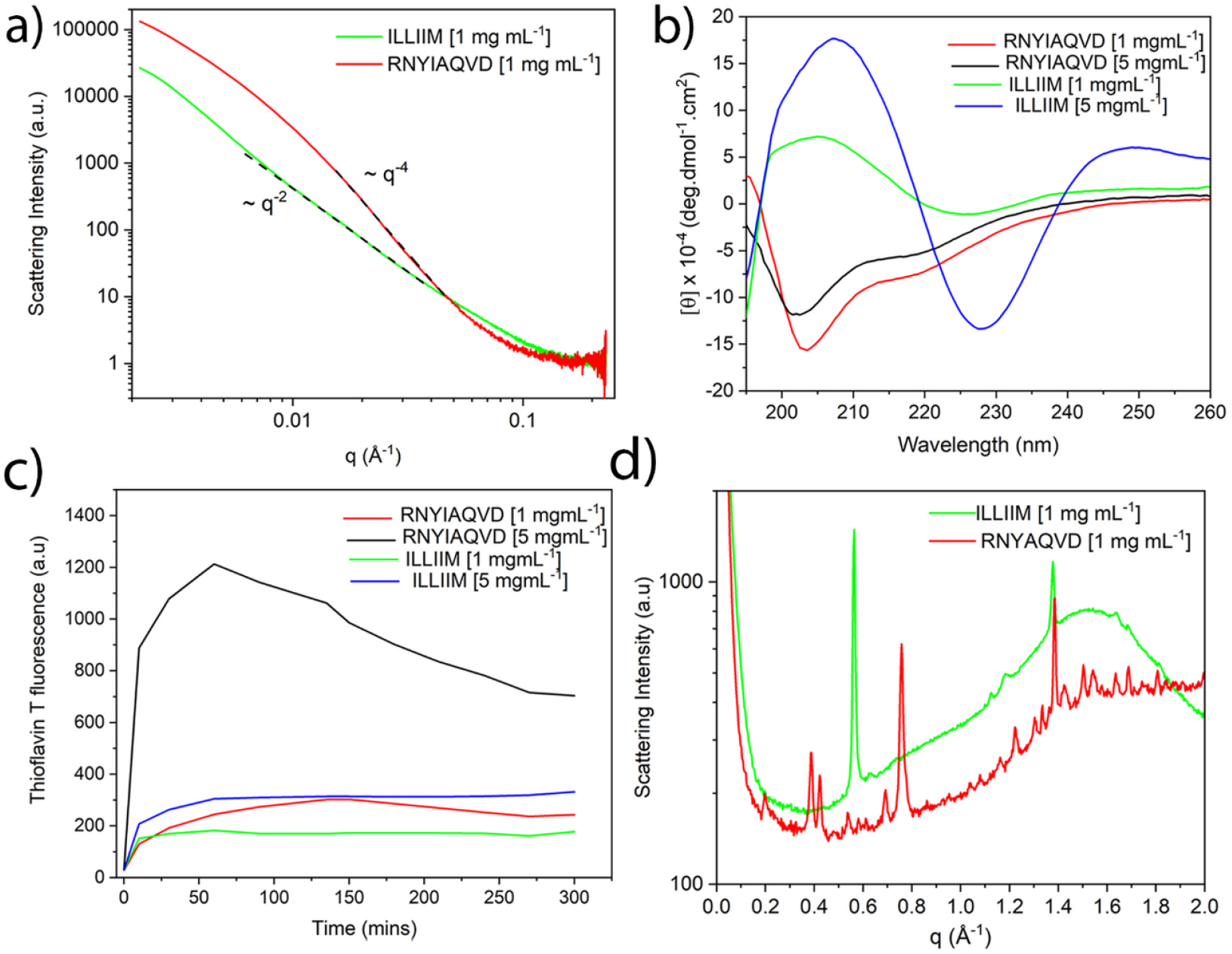
Spectroscopic analysis of the secondary structure of the peptide assemblies. **a)** 1D SAXS plots showing q^−2^ dependence for ILLIIM (flat ribbon) and q^−4^ dependence for RNYIAQVD (spherical particles), **b)** Circular Dichroism spectroscopy of the two peptide assemblies, and **c)** Thioflavin T spectroscopy to quantify the amount of *β*-sheet formation be the two peptide assemblies **d)** WAXS spectra of ILLIIM and RNYIAQVD assemblies. SAXS and WAXS plots are shown as background subtracted against PBS in the same quartz capillary, and averaged from approximately 15 recorded spectra.

To further investigate the predicted secondary structure of both peptide assemblies we employed the secondary structure prediction software BeStSel (figure S8).^45, 58^ As expected from the classic shape of the spectra the predicted secondary structure of ILLIIM is exclusively made up of *β*-sheets ( 49 %) and *β*-turns (51 %). Interestingly, the BeStSel analysis predicts a mixture of right twisted and left twisted *β*-sheets, with a slight preference of left twist (30 % of the total structure) compared to right twist (19 %). RNYIAQVD however, appears to be composed of a more complex mixture of secondary structures, that are still dominated by *β*-sheets and *β*-turns (42 %). The remainder was composed of helical signal (23 %) and further backbone conformation which could not be assigned (35 %). Interestingly almost all of the *β*-sheets in RNYIAQVD are predicted to be right-handed, with only 0.9 % displaying a left handed twist. Part-helical CD spectra do not necessarily imply helical structure, especially considering that a single octapeptide cannot literally be 23% helix (two residues). Backbone conformation as reported by CD correlates through sheet structure to assembled tertiary structure but no single level of organisation exclusively dictates any other, this is especially true in the case of coupling the twist of a peptide strand to the overall twist of the aggregate, which can relax to meet surface and shape-driven constraints through intersheet and interchain as well as intrachain degrees of freedom.^53^

The amyloid nature of the two assemblies is further confirmed by the WAXS spectra (figure 4d) of the peptide assemblies which possessed a number of strongly diffracting Bragg peaks. Both peptides have a clear peak at 1.38 Å^−1^ corresponding to a d-spacing of 4.6 Å which is very indicative of an amyloid assembly composed of extended beta sheets.^59^ ILLIIM also has a very strong Bragg reflection at 0.58 Å^−1^ (11 Å) corresponding to a typical intersheet spacing given moderately bulky hydrophobic sidechains forming a steric zipper. RNYIAQVD has a number of well-defined Bragg peaks in between 0.3-0.8 Å^−1^ which are consistent with a mixture of first and second order reflections corresponding to an amyloid-like 3D symmetry. Discovery of sub-Ångstrom resolution structures from solution WAXS is not at this time a very tractable problem, however given the simple nature of the scattering from the ILLIIM system it was possible to prepare an atomistic model matching the observed peaks. The sheet-like shape factor and the presence of peaks at roughly 2*π*/4.6 Å^−1^ and 2*π*/11 Å^−1^ indicate assembly dominated by the hydrogen bonding axis (with the typical parallel beta sheet period of ≈4.8Å) and by a sidechain-sidechain hydrophobic zipper interface. A candidate structure of size 6×50×1 peptides was constructed following this geometry and found to reproduce the observed WAXS and to fully exclude water at the steric zipper (figure 5).

**Figure 5.**
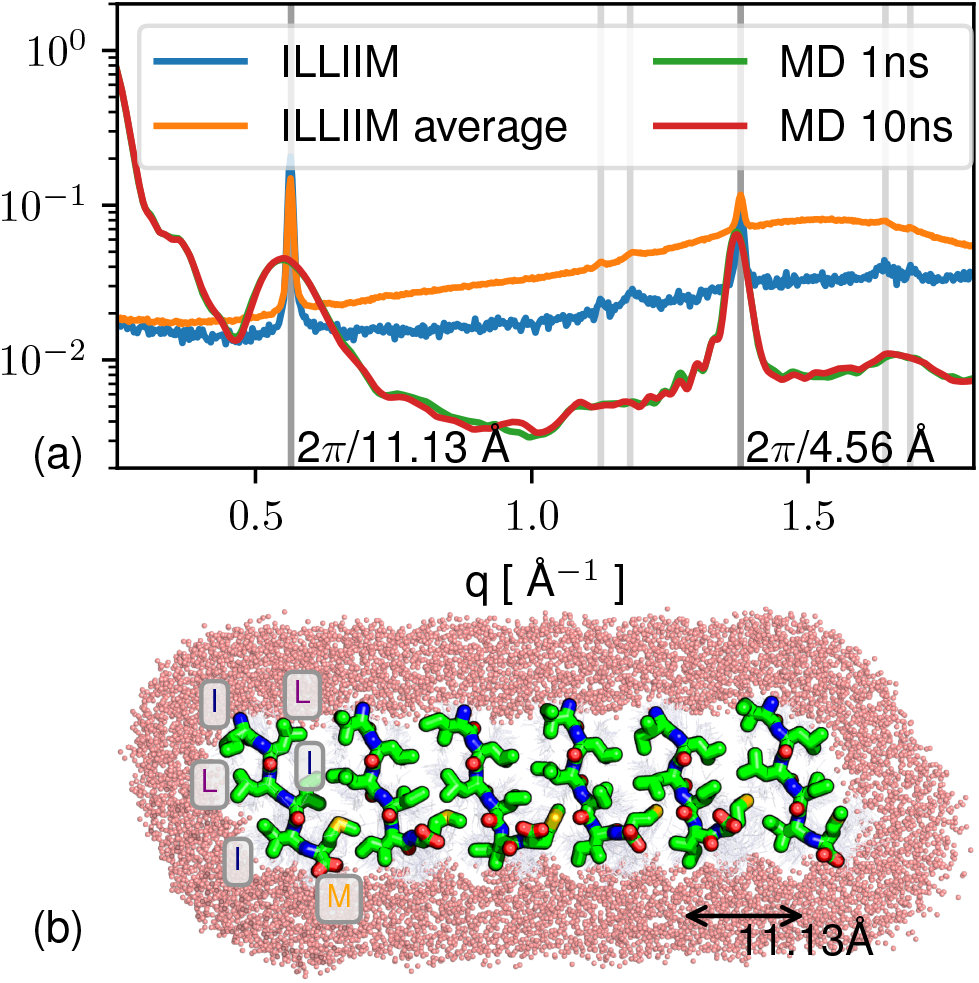
Comparison of scattering peaks between atomistic model 2D amyloid sheet and solution SAXS/WAXS for ILLIIM. **a)** The two major peaks (dark grey) agree well, and some agreement in second order peaks (light grey) is also present. **b)** The modelled 2D structure (parallel *β* hydrogen bonding axis goes into page). Hydrophobic sidechains form a typical amyloid steric zipper. It is not proved that the solution structure is strictly 2D, however the modelling does show that the SAXS/WAXS is consistent with an absence of full 3D symmetry in solution.

The thioflavin T stain, which becomes highly fluorescent upon binding to *β*-sheets present in amyloid fibrils was used to assess the assembly kinetic of both ILLIIM and RNYIAQVD (figure 4c). Both peptides show rapid assembly kinetics reaching a plateau after 30-60 minutes. This behaviour is typical of amyloidogenic short peptides, which can assemble very rapidly without the lag phase typically seen in the amyloid kinetics of longer polypeptide sequences.^53, 60^ This is unsurprising as the short peptides have no additional non-amyloidogenic sequence to reduce the assembly kinetics of the peptides. The ThT signal for ILLIIM at 5 mg mL^−1^ plateaus at about 300 a.u., this is slightly stronger than the maximum signal generated from mature fibrils of the somewhat homologous peptide ILQINS, which was around 250 a.u.^51, 53–55^ suggesting that the amyloidogenicity of the two peptides is comparable. RNYIAQVD, whilst showing similar ThT values at low concentrations (1 mg mL^−1^), generated a ThT signal nearly 3 times as large at 5 mg mL^−1^ suggesting that the assembly of this peptide is highly concentration dependent. For reasons yet unknown, it seems that RNYIAQVD appears to reach a maximum ThT value and then begin to drop, this can be seen at both concentrations but is most obvious at the higher concentration. This could be due to a reversible self-assembly as seen in other functional amyloids^56, 59–61^, or to self-quenching of the amyloid bound aromatic ThT molecules, or simply to a reduction of exposed ThT-binding sites as larger aggregates with a lower surface area to volume ratio come to dominate the solution.

Cytotoxicity of both studied peptides also began to drop slightly (without strong statistical significance) at the highest concentrations tested (*vide infra*, figure 6), together with the drop in ThT response this supports the existence of a kinetically or thermodynamically available aggregate structure with reduced ‘amyloid activity’. This is reminiscent of strongly amyloid-correlated diseases such as AD, where the toxicity of amyloid can vary dramatically, with the relation between the amount of amyloid deposited to the progress of the disease being idiosyncratic and highly non-linear.^62^

**Figure 6.**
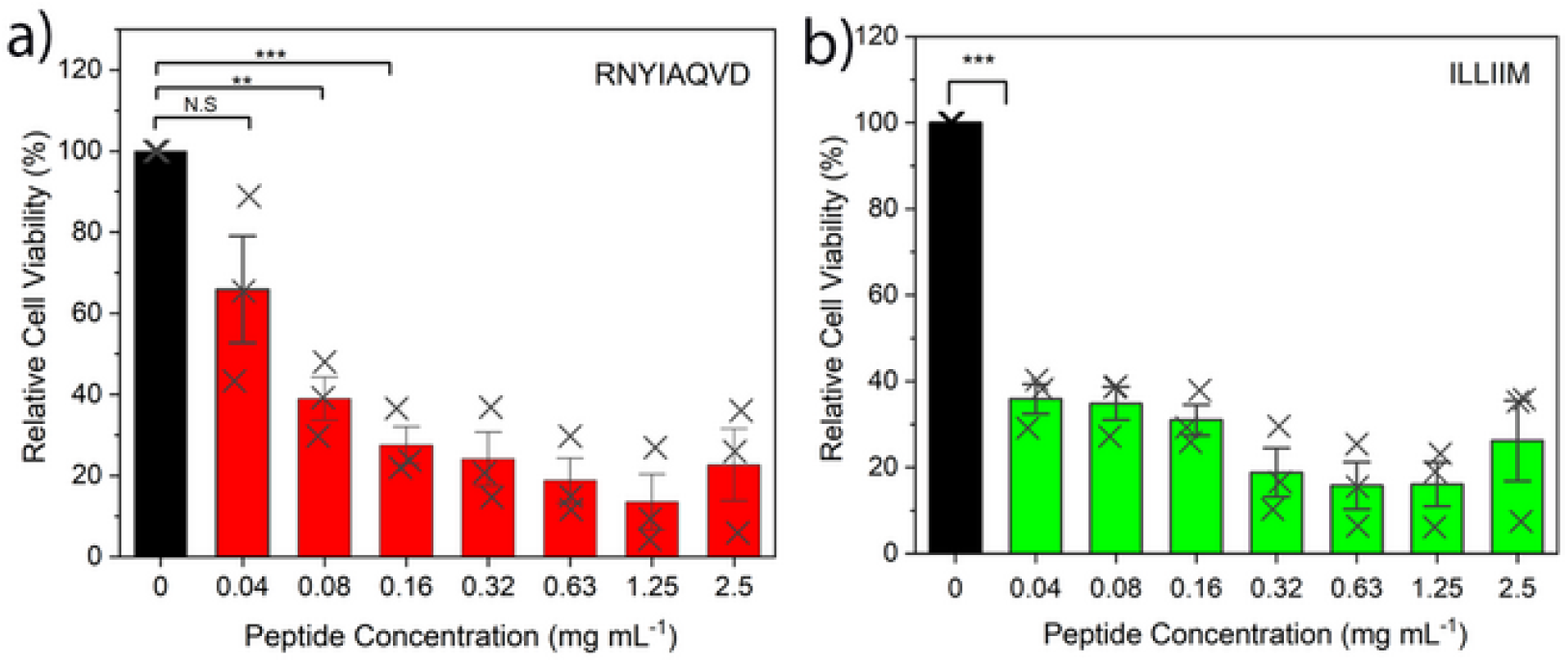
Cytotoxicity assays of RNYIAQVD and ILLIIM assemblies over a range of concentrations. Error bars represent 1 Standard Deviation. Statistical analysis performed by one way ANOVA with Tukey comparison. N.S. = No statistical significance, ** = p < 0.0005, *** = p < 0.0001.

### ILLIIM and RNYIAQVD peptide assemblies are both highly toxic towards the neuroblastoma cell line

Given the physical evidence, and the discussions referred to in the Introduction of various means by which SARS-COV-2 and other viral infections could enhance their fitness (to the detriment of the host) by production of amyloidogenic peptides, we hypothesised that the SARS-COV-2 viral transcript fragments ILLIIM and RNYIAQVD are toxic to human neurons. This is in particular supported by the previously reported neuroinvasive capabilities of of SARS-COV-2,^15, 16^ the noted similarities of the symptoms to a (hopefully transient form of) Alzheimer’s disease^5^ and the previous detection of amyloid assemblies driven by other viruses.^17^ To investigate this further we performed cytotoxicity assays of the two peptide sequences against a human derived neuroblastoma cell line (SH-SY5Y) often used as a model cell line for studying Parkinson’s and other neurodegenerative diseases.^63^ We found that both assemblies were highly toxic after 48 h incubation with the peptide assemblies. Concentrations as low as 0.05 and < 0.04 mM (for RNYIAQVD and ILLIIM respectively) were seen to kill > 50% (IC50) of the cell lines (Figure 6). This toxicity in relation to concentration is similar to that reported for A*β*42^64^ although expression levels and time-envelopes (sudden for COVID versus chronic for AD) are likely to be very different. The toxic nature of these amyloid assemblies warrants further investigations into the potential presence of amyloid aggregates from SARS-CoV-2 in the CNS of COVID-19 patients, and the potential responsibility of amyloid for some of the neurological symptoms observed.

### Concluding Remarks

Using a bioinformatics approach we identified two strongly amyloidogenic subsequences from the ORF6 and ORF10 sections of the SARS-COV-2 proteome. Nanoscale imaging, X-ray scattering, molecular modelling, spectroscopy and kinetic assays revealed that these self-assembled structures are amyloid in nature, and screening against neuronal cells revealed that they are highly toxic (approximately as toxic as the toxic amyloid assemblies in Alzheimer’s disease) in relation to a cell line frequently used as a neurodegenerative diseases model.

The neuroinvasive nature of SARS-COV-2 has been established previously, therefore it is entirely plausible that amyloid assemblies either from these ORF proteins or other viral proteins could be present in in the CNS of COVID-19 patients. The cytotoxicity and protease resistant structure of these assemblies may well explain the lasting neurological symptoms of COVID-19, especially those which are novel in relation to other post-viral syndromes such as that following the original SARS-COV. The outlook in relation to triggering of progressive neurodegenerative disease remains uncertain, given the typically slow progress of neurodegenerative disease then if such a phenomenon exists it will most probably take some time to become evident epidemiologically.

## Acknowledgements

NPR would like to acknowledge The La Trobe Institute of Molecular Sciences (LIMS) for the receipt of a Nicholas Hoogenraad fellowship. Dr. Susi Seibt for assistance on the SAXS/WAXS beamline at the Australian Synchrotron. This research was undertaken, in part, on the SAXS/WAXS beamline at the Australian Synchrotron, part of ANSTO. Molecular dynamics calculations made use of the HPC service of the University of Luxembourg^65^.

## Notes

### Competing Interest Statement

The authors have declared no competing interest.

