## Supplementary Figures for "Neurotoxic Amyloidogenic Peptides Identified in the Proteome of SARS-COV2: Potential Implications for Neurological Symptoms in COVID-19"

***Supplementary Data***

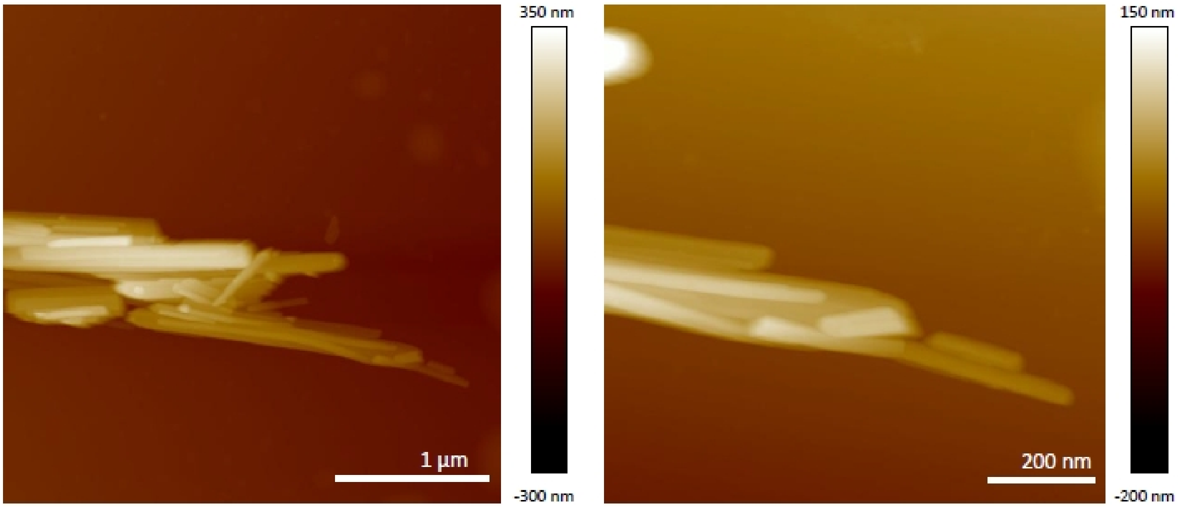


Figure S1: AFM Images of RNYIAQVD assemblies [1 mg mL$^{-1}$] after 2 h assembly

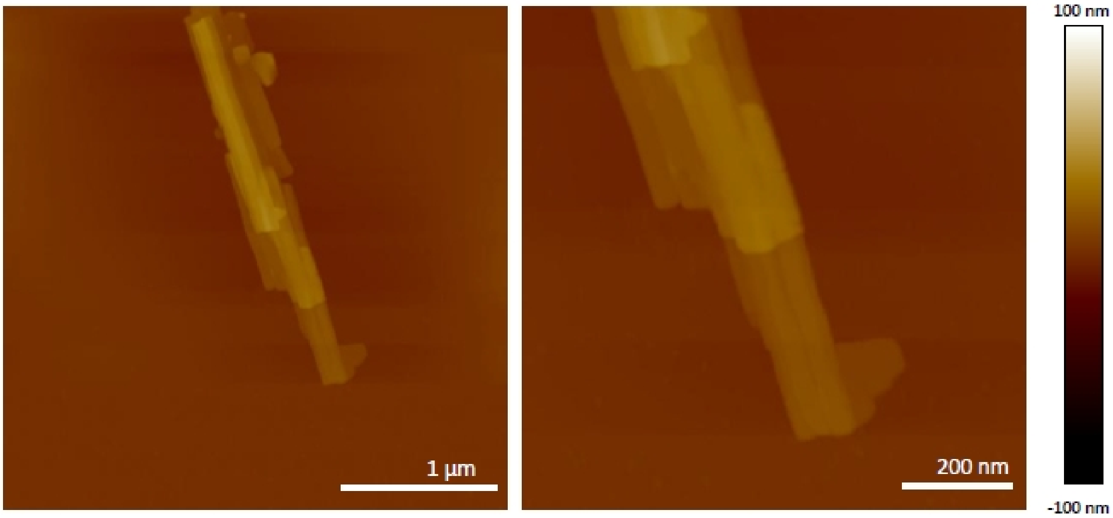


Figure S2: AFM Image of ILLIIM assemblies [1 mg mL$^{-1}$] after 2 h assembly

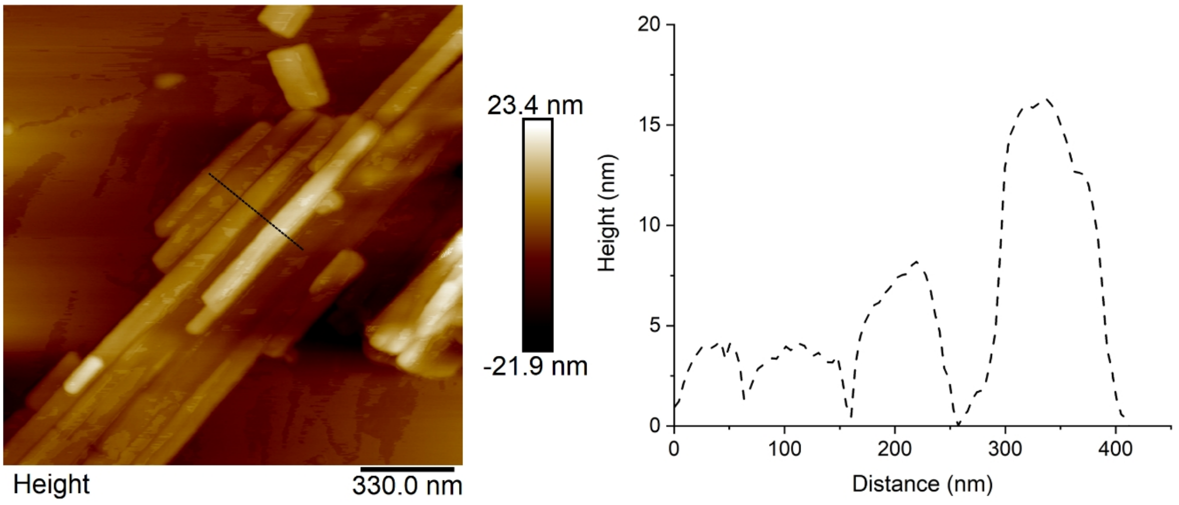


Figure S3: Line scans of ILLIIM assemblies (5 mg $mL^{-1}$)

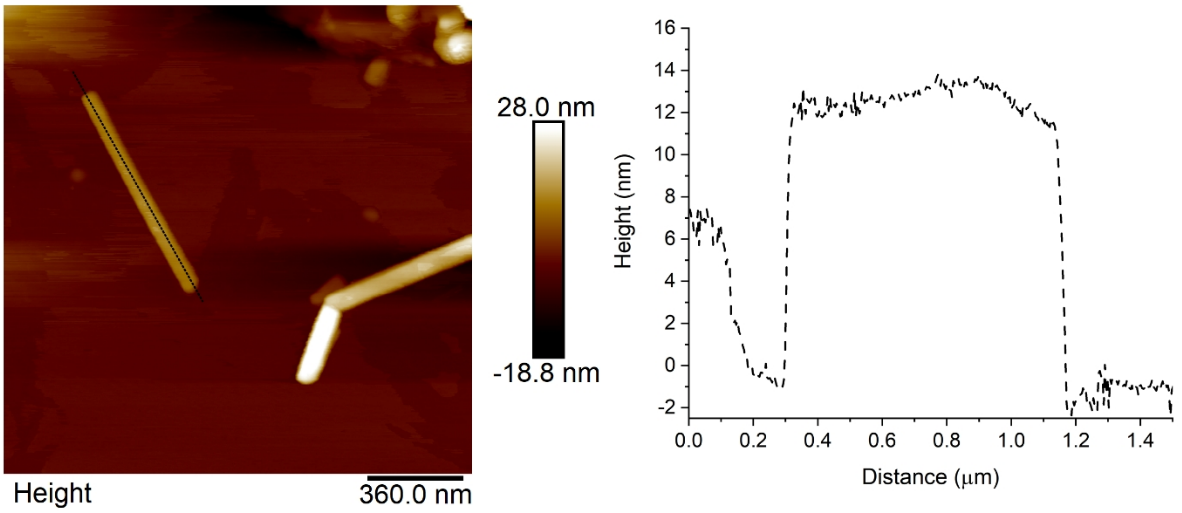


Figure S4: Line scans of ILLIIM assemblies (5 mg $mL^{-1}$)

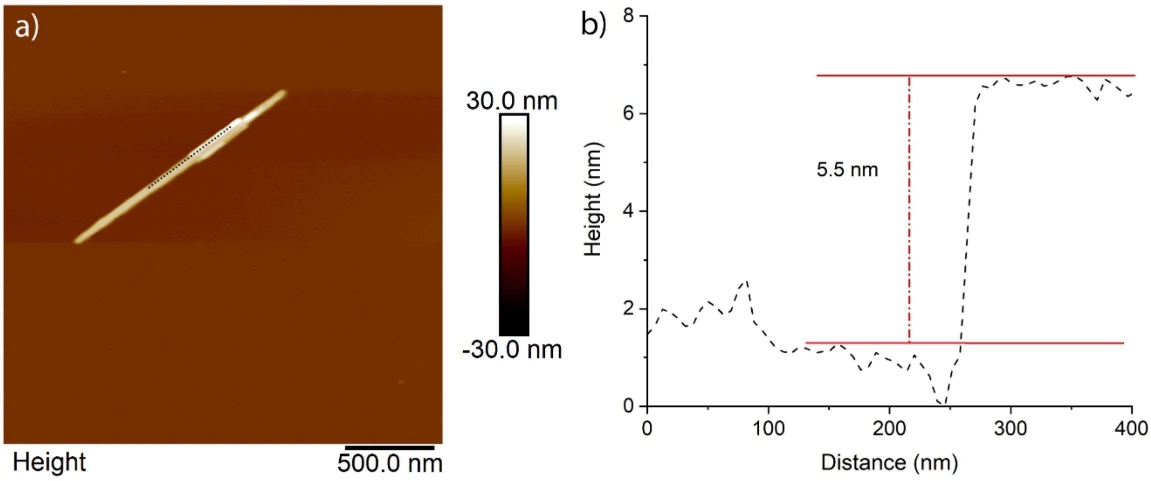


Figure S5: Line scans of RNYIAQVD assemblies (5 mg mL$^{-1}$)

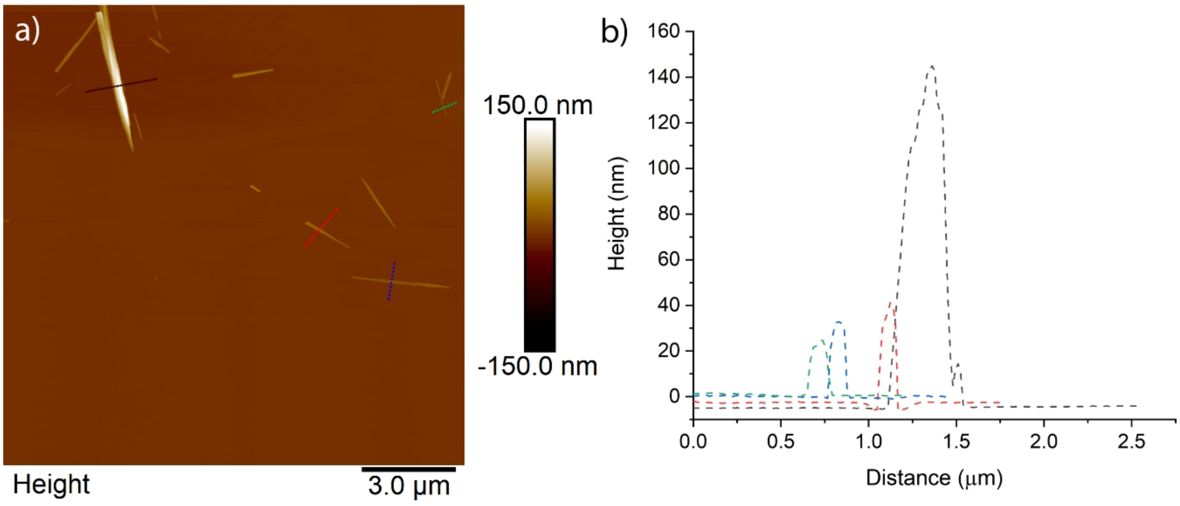


Figure S6: Line scans of RNYIAQVD assemblies (5 mg mL$^{-1}$)

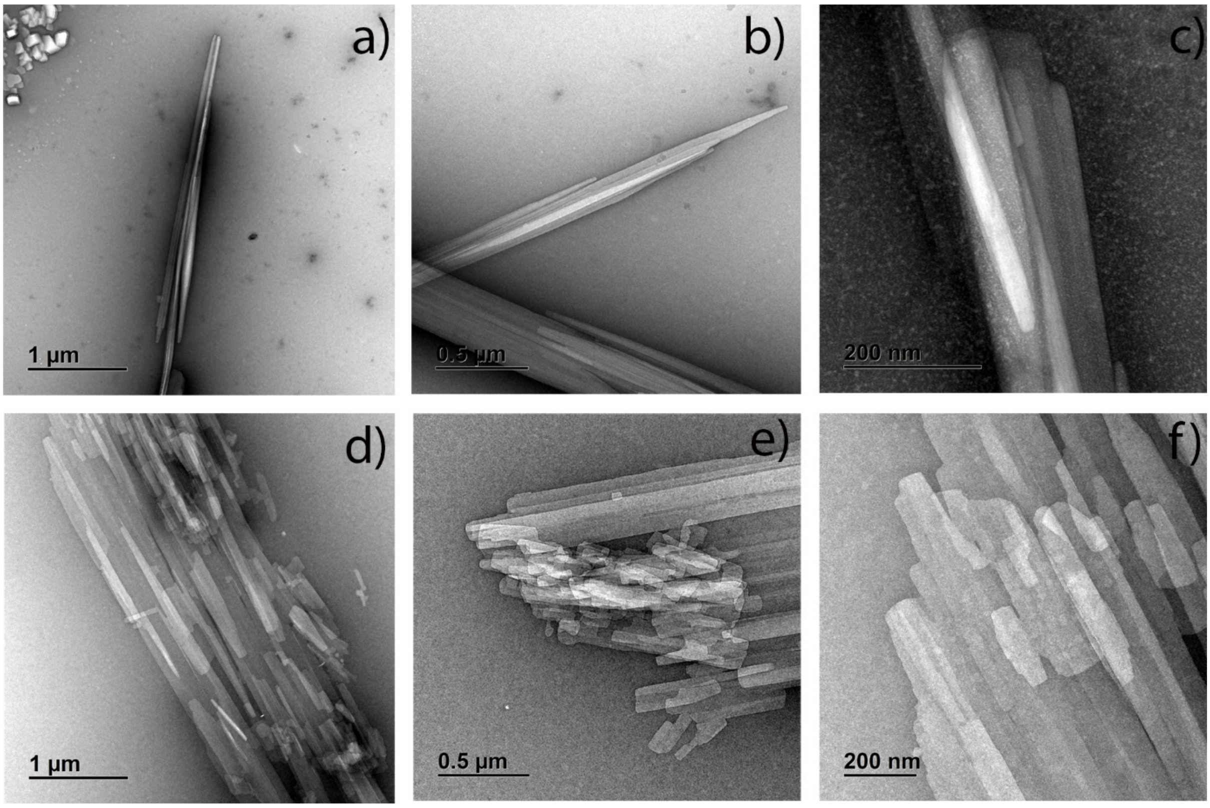

Figure S7: Additional TEM images a-c) RNYIAQVD d-f) ILLIIM

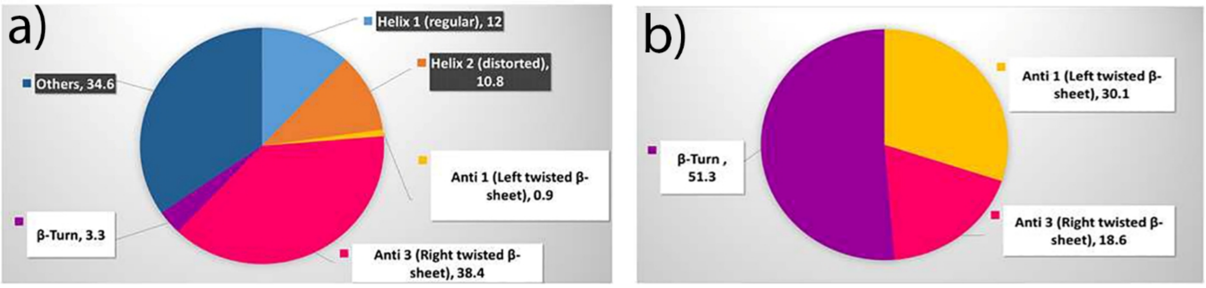


Figure S8: Secondary Structure Analysis from BestSel of a) RNYIAQVD and b) ILLIIM

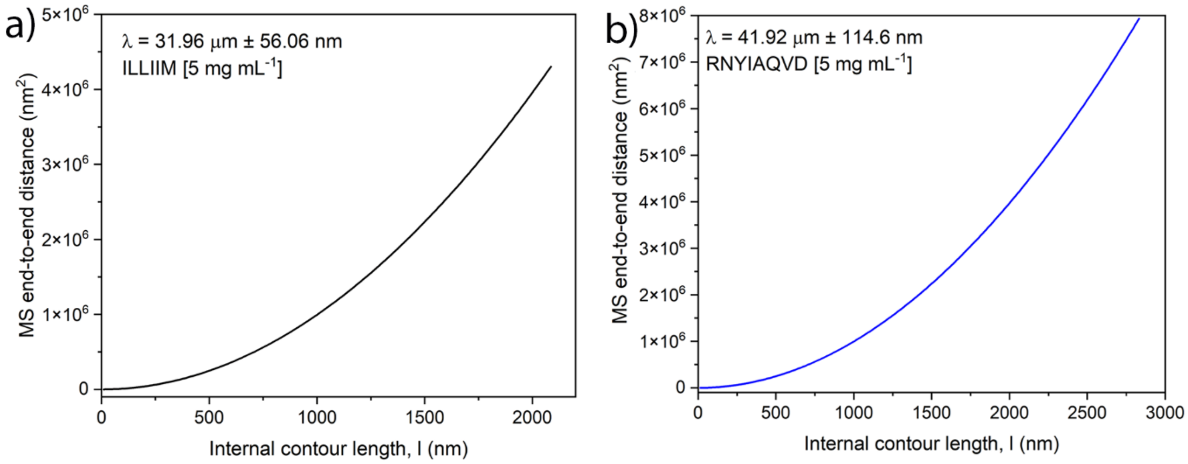


Figure S9: Mean-Square end-to-end persistance length calculations for both peptide assemblies

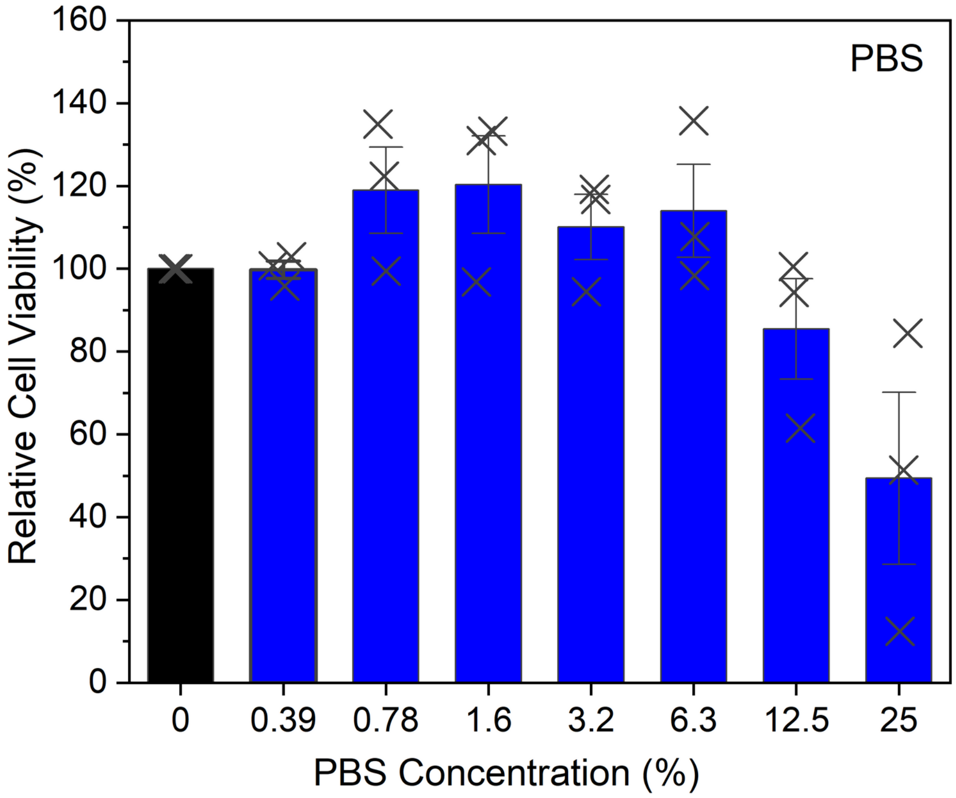


Figure S10: Cytotoxicity assays showing the toxicity of PBS in the concentrations used for peptide cytotoxicity  assays
